# The difference in FRET efficiency fluctuation at each end of the actin filament

**DOI:** 10.1101/2022.07.15.500159

**Authors:** Takahiro Mitani, Ryota Mashiko, Miku Nezasa, Ichiro Nishikata, Kenji Kamimura, Hajime Honda, Ikuko Fujiwara

**Affiliations:** Department of Bioengineering, Nagaoka University of Technology, Niigata, Japan; Department of Materials Science and Bioengineering, Nagaoka University of Technology, Niigata, Japan; Deptartment of Electronic Control Engineering, NIT, Nagaoka College, Niigata, Japan; Electrical & Mechanical Systems Engineering Advanced Course, NIT, Nagaoka College, Niigata, Japan

**Keywords:** actin, fluctuation, FRET, filament asymmetry, direct observation

## Abstract

Actin filaments are involved in various cell motility processes. Actin polymerisation is primarily governed by monomer association and dissociation occurring at the rapid-growing end called the barbed end, which generates the force to push the plasma membrane forward. Individual actin filaments bind to one nucleotide and its hydrolysis energy is used to maintain the filamentous form by changing the characteristics of the subunits. The asymmetry of the individual actin filaments is also important for detecting the asymmetry of the cell. However, asymmetry at the subunit level including conformational and temporal changes in actin filaments has not been visualized yet. Here, we used “Förster (or fluorescence) resonance energy transfer (FRET)-actin filament” by copolymerising an equal amount of donor and acceptor labelled actin. FRET efficiency change was measured along each actin filament under a light microscope. The FRET efficiency was lower near the end region than in the interior regions. Fluctuations in the FRET efficiency (*f*_FRET_) were used to monitor local flexibilities along each actin filament. The *f*_FRET_ was larger near the end region than the interior region. Our quantified data showed that spatial change of *f*_FRET_ along actin filaments was rapidly decayed from the barbed end from near the pointed end toward internal region, suggesting that the behaviour of actin subunits near ends is affected from each end. Our result revealed that actin filaments have different orientations locally. These orientations appear when actin forms filaments, which may contribute the cell motility.

## 1 Introduction

Actin polymerisation and depolymerisation dynamics are essential for cell motility. Under physiological conditions, polymerisation mainly occurs at the barbed end with ∼18 times tighter binding affinity than the other end called the pointed end. Each actin molecule binds to the ends of the actin filaments through their intrinsic binding sites. Consequently, the actin subunits align and form a filament facing the same direction. Thus, for filament polarity, the binding sites linking each actin subunit are crucial; though this alignment might occasionally fail when the filament size is very small (Holmes et al., 1990; Kabsch et al., 1990; Horan et al., 2020). Each actin can bind and hydrolyse ATP as the polymerisation progresses. The incorporated actin subunits contain ADP-Pi-actin, which is the most stable form (Fujiwara et al., 2007). With the slow dissociation of ɤ-phosphate (Pi) (Blanchoin and Pollard, 1999), the bound nucleotide becomes ADP, and the actin subunit dissociates from the terminal end of the filaments, a process known as depolymerisation. ATP hydrolysis related to (Blanchoin and Pollard, 1999) polymerisation dynamics is often expressed as “aging factors” of actin filaments. Actin aging alters filament asymmetry and properties such as torsional and bending rigidity, bundling, and severing (Pollard, 1984; Carlier et al., 1986; Kinosian et al., 1993; Winkelman et al., 2016; Bibeau et al., 2021). The actin-binding protein ADF/cofilin senses the asymmetry of actin filaments via ATP hydrolysis. It binds cooperatively to the side of actin filaments, with a change in binding affinity based on the nucleotide state and tension of the actin filaments (Hayakawa et al., 2011; Bibeau et al., 2021). Besides, the ADF/cofilin-bound actin subunit altered the conformation of the C-form of the actin subunit (Tanaka et al., 2018). Therefore, understanding the asymmetry of actin filaments is a crucial regulatory mechanism for the actin cytoskeleton. However, the status of actin subunits in each actin filament is not fully understood, because the dynamic behaviour of actin subunits in the filament is not easy to visualise in real-time at a single filament level. Instead, recent structural biology studies using cryoEM, crystal structure, all-atom simulations, and molecular dynamics calculations have unveiled the molecular status of actin subunits in filaments related to ATP hydrolysis. CryoEM studies have revealed that the domain binding and canonical actin interactions, especially the multiple interactions of subdomain-3, are the key to the binding of subdomain-2 of the adjacent actin subunit (Galkin et al., 2015; Chou and Pollard, 2019). Oda et. al. demonstrated by cluster analysis using rigid bodies of the actin structure, that the structure of actin in the filament fluctuates with greater fluctuations observed in the monomeric actin (G-actin) structure (Oda et al., 2019). All-atom simulations suggest that the conformational fluctuation of ∼100 actin subunits from the terminal ends is larger than the interior actin subunits (Zsolnay et al., 2020). The authors described that the pointed end subunits were very rigid concerning each other in the ADP state, in contrast to the loosely attached and flexible ATP-bound barbed ends. Although this is an important finding for understanding the dynamics of actin at local positions along the actin filament, the source data is derived from structural information, which possibly misses data from temporal information.

Actin filament aging was visualized using Total Internal Reflection Fluorescence (TIRF) microscope (Jégou et al., 2011; Fujiwara et al., 2018). The depolymerization process was initiated by washing the free actin monomers. Slow depolymerization, corresponding to ADP-Pi bound actin subunits, was followed by fast depolymerization, corresponding to ADP-bound actin subunits. These results confirmed that due to ATP hydrolysis, the terminal actin subunits tend to bind ADP-Pi and the interior actin subunits bind ADP. These researches strongly suggest that the conformation of an actin filament is not same. Rather, dozens of actin subunits at each terminal end show different dynamics from the interior actin subunits. In this study, we investigated whether there is a difference in conformational fluctuations along actin filaments by monitoring fluorescence fluctuations.

FRET is a potent approach for measuring the distance between the donor and acceptor using fluorescence intensity, enabling us to trace conformational changes at the single-molecule level (Ha et al., 2002; Kozuka et al., 2006; Lam et al., 2010; Santoso et al., 2010; Morimatsu et al., 2012; Noguchi et al., 2015). When the distance between the donor and acceptor is sufficiently long, the donor fluorescent signal can be detected. Whereas, the donor signal becomes low when the distance is short, because the fluorescent energy of the donor is transferred to the acceptor. FRET efficiency can be inversely proportional to the distance. FRET efficiency is largely observed in the distance range of 3–8 nm and is inversely proportional to the sixth power of distance (Kozuka et al., 2006; Hirata and Kiyokawa, 2016). Here, we observed that FRET-actin filaments were formed by mixing Dylight 550- and Dylight 650-labeled actins, and the FRET fluctuation occurring in one actin filament was measured under a fluorescence microscope. This approach enabled us to obtain spatiotemporal information on the FRET efficiency. The fluctuation in FRET efficiency (*f*_*FRET*_) was larger at both ends than in the interior of the actin filaments and decreased as it approached the interior regions. At the barbed ends, the decay was faster than at the pointed ends, although the magnitude of the fluctuation was larger at the pointed ends. Our results suggest that actin regulators sense differences in molecular dynamics at both ends of actin filaments, which may alter the kinetics.

## 2 Materials and methods

### 2.1 Proteins

Actin was eluted from acetone powder of rabbit or chicken skeletal muscle with G-buffer (2 mM Tris-HCl, pH 8.0, 0.2 mM ATP, 0.1 mM CaCl_2_) in the presence of 1 mM DTT based on the previous literature (Spudich and Watt, 1971)Spudich and Watt, 1971. After acetone powder was removed from the eluate with 3-fold of filter paper (No.1 (Avantic, Japan)) attached to aspiration at 4 °C, contaminations were removed by ultracentrifugation (223,000 × *g* rpm for 40 min at 4 °C). The supernatant was polymerised by adding the final 1 mM ATP, 1 mM MgCl_2_ and 50 mM KCl and incubated at 25 °C for 1 h. The Polymerised actin was pelleted by ultracentrifugation (223,000 × *g* at 25 °C for 1 h), resuspended with G-buffer with 1 mM DL-Dithiothreitol (DTT), and dialysed in G-buffer overnight on ice. Large contaminations and non-depolymerized actin were removed by ultracentrifugation (40,000 rpm at 4 °C for 1 h). Monomeric actin (G-actin) was loaded onto a 1 × 7 cm desalting column (Toyopeal 40S-HW, TOSOH, Japan) equilibrated with 1 mM ATP pH 7.0 and 0.1 mM CaCl_2_. The peak fraction was pooled and the concentration of purified actin was determined by differences in extinction coefficients between 310 and 290 nm, namely 0.62 (ε_290_-ε_310_ = 0.62).

G-actin was rapidly and well mixed with 9.2 mM of Dylight 550-maleimide or Dylight 650-maleimide (Thermo Scientific, United States) that were dissolved in dimethyl sulfoxide for a few minutes (to be an equal molar ratio) subsequently mixed and incubated at 4 °C overnight in dark. The labelling process was stopped by the addition of 28 mM 2-mercaptoethanol. The concentration of labelled actin was measured at 290 nm and 557 or 655 nm was used for absorbance of Dylight 550 or Dylight 650 with molar extinction coefficients of 150,000 or 250,000. The ratio of labelled actin to total actin was calculated using correction factors of 0.081 for Dylight 550 and 0.037 for Dylight 650.

Biotinylated actin was prepared by mixing G-actin and biotin-PE-maleimide (Dojindo, Japan). Biotin was dissolved in dimethyl sulfoxide (DMSO), and added in an equal molar ratio to actin, and incubated overnight at 4 °C using an end-end mixer. The free biotin was removed using a column (PD-25, Cytiva, Japan). The column was equilibrated with 1 mM ATP pH 7.0 and 0.1 mM CaCl_2_ before loading labelled actin. The concentration of biotinylated actin was the same as that of actin by considering that all actin was biotinylated.

To prepare for FRET conditions, 24 μM actin stock, 30% Dylight 550-maleimide-actin, 30% Dy650-maleimide-actin, 1% biotinylated actin, and 39% non-labelled actin were mixed. Afterward, the 5× polymerisation buffer (final10 mM Tris-HCl, pH 8.0, 100 mM KCl, 2 mM ATP, 1 mM MgCl_2_, and 0.1 mM EGTA, pH 9.1) was added and incubated overnight at 4 °C. Phalloidin-bound actin was prepared by adding 2 mM phalloidin to 24 μM polymerised actin for a 1:1 molar ratio and incubated overnight in dark at 4 °C. Anti-biotin antibody, gelsolin from bovine plasma, goat anti-gelsolin antibody, anti-Rat IgG were purchased from Sigma Aldrich, Japan.

### 2.2 Glass Preparation

The glasses for the observation chambers were prepared as described previously (Ozawa et al., 2019). Briefly, two different sizes of coverslips, 18×18 and 25×50 mm (Matsunami, Neo No. 1), were soaked in piranha solution containing a 7:3 (v/v) mixture of sulfuric acid (H_2_SO_4_) and 30% hydrogen peroxide (H_2_O_2_) for 30 min. Both coverslip sizes were rinsed with distilled water and then soaked in a 5% TMCS-ethanol solution (mixture of 5% TMCS and 95% ethanol) overnight at room temperature. The coverslips were transferred and stored in absolute ethanol in glass jars with a lid and used within two days. The coverslips were rinsed with two cycles of absolute ethanol and pure water and later dried in clean air before use.

### 2.3 Flow chamber preparation

Flow chambers were prepared daily, as per the procedure as described previously (Kron and Spudich, 1986). The flow chamber was prepared by mounting a small coverslip on the larger coverslip placing two double-sided tapes of 0.15 mm thickness (ST415 25 × 30, 3M, Japan) in parallel, leaving a gap of ∼5 mm. The size of the flow chamber was approximately 18 × 5 × 0.15 mm with both sides open (∼10 μl volume).

### 2.4 Specimens

The anti-biotin antibody (5 μg/ml) was loaded directly into the chamber via capillary action. The unbound anti-biotin antibody was washed out with a chamber volume of the 5×polymerisation buffer at one end and subsequently absorbed from the other side using a piece of coffee filter paper. Polymerised actin was diluted to 5 nM with 1× polymerisation buffer, containing 10 mM ascorbic acid as an oxygen scavenger, immediately before loading into the chamber. Unbound actin filaments were washed out with polymerisation buffer containing 10 mM ascorbic acid. FRET signals were observed immediately. To prevent fluid flow onto the top of the coverslip due to the electrostatic effect caused by the hydrophobic surface of the coverslip, occasionally sub-microliters of dH_2_O were dropped before the coverslip was mounted with a double-sided tape. Two approaches were utilized to identify the barbed end. The first step was to elongate actin at the barbed end with 30% Dylight 650-labelled actin. The second approach was used to identify the barbed end. Gelsolin, a known actin severing/barbed-end capping protein (10 nM), was mixed with 66 μM FRET actin filaments in a sample tube. Gelsolin on the barbed end was visualised by decollating the goat anti-gelsolin antibody with two concentrations of anti-rat fluorescent secondary antibody.

### 2.5 Observation

The chambers were mounted on an Olympus IX-71 microscope with a 100× objective lens (Olympus, Plan Apo, Japan). White light from an LED lamp (Lumencor, SOLA, United States) was passed through a filter cube f containing an excitation filter (FF01-542/27-25), dichroic mirror (FF560/659-Di01-25×36), and emission filter (FF01-577/691-25). To observe the Dylight650 actin filaments, a filter cube containing HQ620/60X 62032, FF650-Di01-25×36, and HQ700/75M 57563 was used. The final power of the excitation light was attenuated using neutral density filters of up to 5 mW. All filters were purchased from Semrock, United States Images were recorded using an EM-CCD camera (iXonX3, Andor Technology, United States) attached to the side port of the microscope through a double view (U-SIP, Olympus, Japan).

### 2.6 Image analysis

Images were recorded in uncompressed AVI format (0.1-sec exposure). We used an algorithm in MATLAB (MathWorks, United States) to measure the fluorescence fluctuation along each actin filament. After manually setting the polygonal line along the actin filaments containing 2 μm extrapolation, the Resion of Interest (ROI) (Φ = 1.5 μm) automatically traced the line. The sum of the fluorescent intensity of the ROI was used as each data point. The ROI position was shifted every 0.1 μm (∼1 pix) along each actin filament (including oversampling) to avoid missing local fluorescent fluctuations. To subtract the background in each image, an identical trace was performed 2-3 μm apart from the targeted actin filaments where no actin was present. FRET efficiency was determined from the ratio of acceptor (Dylight 650) / donor (Dylight 550) intensity. The *f*_*FRET*_ for 30 s recording were determined from temporal changes in FRET efficiency from a single and identical ROI on actin filaments taken every 0.1 sec.

## 3 Results

### 3.1 FRET signal is measured from an actin filament

We measured the fluctuations in FRET signals from subunits of actin filaments. FRET actin filaments were prepared by polymerising a mixture of Dylight 550 labelled actin (30%) and Dylight 650 labelled actin (30%), and unlabelled actin (39%). The either of dyes was conjugated to 374-cysteine residue on actin. To anchor FRET actin filaments on the glass surface coated with a 5.0 μM anti-biotin antibody, 1% biotinylated actin was also added (Figure 1A) to the FRET actin filament. After polymerising actin in the dark at concentrations over 66 μM in a sample tube overnight at 25 °C, polymerised FRET actin was diluted to 5 nM with the observation buffer and loaded into the glass chamber using spontaneous capillary flow. The anchored actin filaments did not show any detectable wobbling movement during the 30 s observation period (Figure 1B), indicating that the FRET-actin filament was fixed on the glass surface. Fluorescence images of both the donor and acceptor from the same filament were recorded simultaneously using a dual-view microscope (Figure 1A). During the observation, every 1/30 s for 30 s, the apparent length of actin filaments did not change, probably because the ratio of the fluorescently labelled actin was high in this assay at 60%. Previous reports have shown that the rate of actin polymerisation and depolymerisation decrease with an increasing ratio of the labelled actin (Amann and Pollard, 2001; Kuhn and Pollard, 2005; Fujiwara et al., 2018). The analysis was simplified based on the fact that depolymerisation was negligible in this observation time frame (Figure 1B). Next, to confirm that the fluorescence from the acceptor dye resulted from the energy transferred from the donor fluorescence, we adopted two essentially different approaches. The first approach was acceptor photobleaching. Excess light at 650 nm was irradiated to bleach the acceptor dyes, followed by normal observation of donor illumination (as shown in Figure 1C, blue decayed plot). After the photobleaching of the acceptor dye, the fluorescence intensities from the donor were increased (as shown in Figure 1C, red plot), indicating that the donor fluorescence is no more transferred to excite the acceptor because the acceptor dyes were no more active. The second approach was performed to test the complementarity of the donor and acceptor fluorescence. Both fluorescence intensities decreased over time (Figure 1D). We corrected both signals using a single exponential decay function to avoid the effects of photobleaching and subtracted 5% of donor intensities from the acceptor due to the transmission characteristics of the filter sets. The intensity profile became in reverse phase after the curves were smoothed by the moving average method for visual clarification (Figure 1D inset), indicating that the acceptor fluorescence was excited using the energy transferred from the donor. Both approaches confirmed that our microscopy settings detect FRET within a single actin filament, and the calculated FRET efficiencies become a parameter to indicate the mobilities of subunits in the filament.

**Figure 1.**
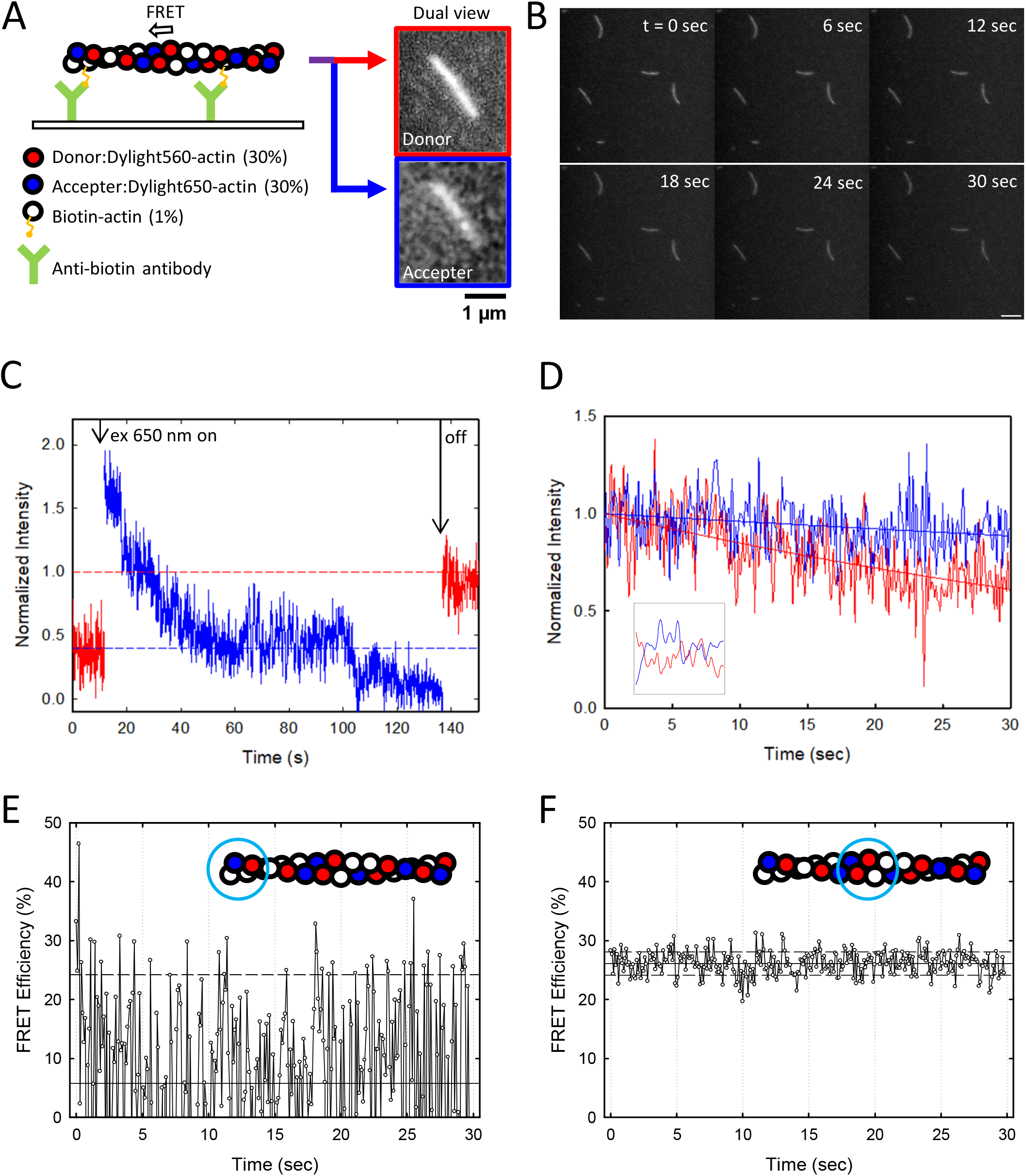
Experimental setup and FRET efficiency on single actin filaments. (A) Schematic illustration to observe FRET actin filaments. Glass surface is coated with anti-biotin antibody (green). 1% BSA was subsequently loaded to block non-specific binding to the glass surface. Actin filaments were diluted to 5 nM with the observation buffer (10 mM Tris-HCl; pH 8.2), 100 mM KCl, 2 mM CaCl_2_, 0.2 mM ATP, and oxygen scavenger), right before loading into the glass chamber. Actin filaments were polymerized by mixing 30% of Dylight-550 (red), 30% of Dylight 650 (blue), and 1% of biotinylated actin (yellow) one hour before use. FRET signals were split with a 561 nm filter and simultaneously captured using a dual-view system. The right panels show a typical actin filament observed under a FRET microscope monitored by Dylight 550 channel (top) and Dylight 650 channel (bottom). (B) No wobbling and length change of actin filaments were detected under this setup for 30 seconds of observation. Still, images show an actin filament every six seconds with Dylight 550 (donor) signal. (C) Fluorescence intensity recovery by the acceptor photo-bleach over time. Signals of the donor (Dylight 550, red) and acceptor (Dylight 650, blue) were normalised and plotted over time. The acceptor signal was forcibly photo-bleached with 650 nm of light during the time frame indicated by arrows. Donor signal intensity increased after the acceptor signal was photobleached, indicating that no acceptor energy was transferred to the donor. (D) Spontaneous photo-bleaching of the donor (red) and acceptor (blue) were measured for 30 seconds and fitted with single exponential decay curves. Inset indicates high throughput intensity changes of the donor and acceptor for 10 seconds. The fluorescence of the acceptor was weak and the donor signal was bright. (E and F) Time course FRET efficiency at the end region (E) and the interior region of an actin filament, as the measured location was circled on the illustrated actin filaments.

Since energy transfer is generated from Dylight 550-actin to Dylight 650-actin in a filament, our data monitored the change at the subunit-subunit level and not within an actin molecule. By labelling one actin with two fluorophores, it is found that the molecular fluctuation of individual actin subunits in a filament takes two states and the transition rate is 0.24–0.34 /s (Kozuka et al., 2006). This rate is 10-fold slower than the sampling rate in our observation, 0.03 (= 1/30) s. Thus, the fast FRET signal from this assay was most likely due to the conformational change caused by the fluctuation of actin subunits in the filament. During the 30 s of observation, FRET efficiency at the end region of the actin filament was constantly low, while it was fluctuating (Figure 1E). This indicates that the location of actin subunits was far each other and not stable. In contrast, FRET efficiency was temporally stable and high in the interior regions of the filament (Figure 1F), indicating that the actin subunits in the interior region are closely and stable located.

### 3.2 Subunit fluctuation near the terminal ends is larger than the interior region of the actin filaments

In previous section, FRET efficiency fluctuated temporally, which arises a hypothesis that it may also differ spatially in actin filaments. Thus, we investigated the configuration between the neighbouring subunits. We scanned FRET efficiency along actin filaments at every 0.1 μm and the average and standard deviation were estimated from 30 s of observation (Figure 2A). This fluctuation in FRET efficiency (*f*_FRET_) was quantified using the standard deviation of FRET efficiencies for 4 s, obtained as the subunit fluctuation at each local position of the actin filament (Figure 2B). Remarkably, the spatial gradient of *f*_FRET_ at one end decreased dramatically towards the interiors of the actin filament, while it decreased gradually at the other end. From this perspective, we plotted the absolute spatial gradient of *f*_FRET_ at 0.2 μm from each end (*Δf*_FRET_, Figures 2C and 2D). One hundred filaments were subjected to *Δf*_FRET_ without distinguishing the end and the distributions (black histograms in Figure 2C). Using the global fitting method, it was found that *Δf*_FRET_ consists of two Gaussian distributions which have proven to be significantly different (P < 10^−19^) between each peak. To investigate whether these distributions originated from the structural polarity of the filament, the *Δf*_FRET_ values from both ends were plotted separately for each actin filament. Our analysis revealed that one end of the actin filament showed a larger *Δf*_FRET_ (Figure 2C, red) than the other (Figure 2C, blue). Even filaments associated with phalloidin had the same properties, the two *Δf*_FRET_ distributions had a statistically difference (P < 10^−4^, Figure 2D).

**Figure 2.**
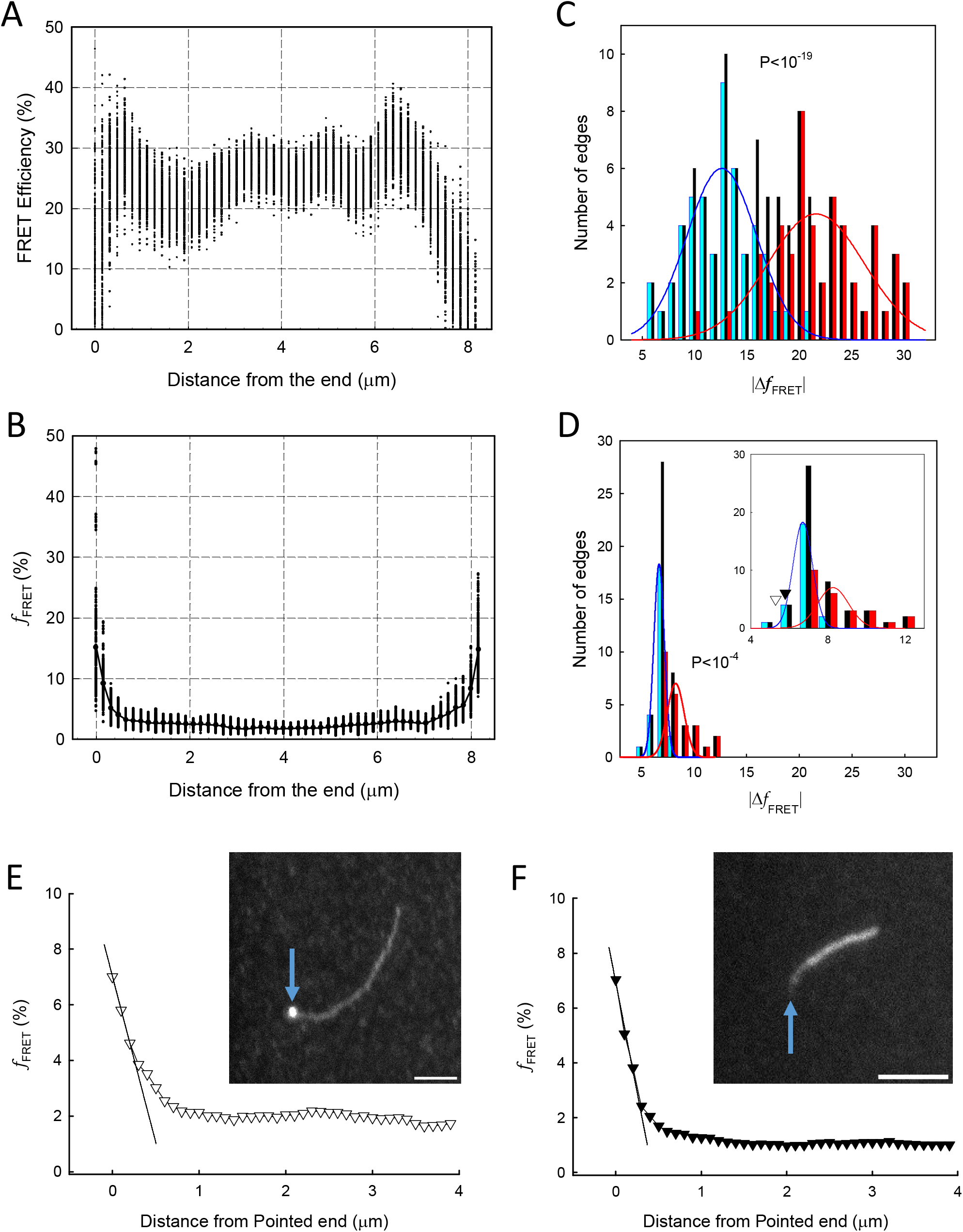
Evaluation of *f*_*FRET*_ and *Δf*_*FRET*_. (A and B) FRET efficiency (A) and fluctuations of FRET efficiency (*f*_*FRET*,_) (B) are plotted along an actin filament at distances of 0.1 μm over 30 seconds of observation. (C and D) *Δf*_*FRET*_ in the regions of 0.3 μm from both ends without (C) and with phalloidin (D). The *Δf*_*FRET*_ from both ends of actin filaments (scrambled) was plotted (black, n = 100). A small *Δf*_*FRET*_ from one end of the actin filament (blue, n = 50) and a *Δf*_*FRET*_ from the other end (red, n = 50) were plotted. Colour Gaussian curves are fitted by using the scramble distribution, the peaks of small and large gaussian. Inset in D shows the magnified image of D. Open and filled triangles show the average *Δf*_*FRET*_ estimated from linear fittings in panels E and F, respectively (average of 10 samples, each). (E and F) The *f*_*FRET*_ from the pointed end of the actin filament. The insets show actin filaments that barbed ends were distinguished by gelsolin (E), which contained anti-gelsolin antibodies decorated with secondary fluorescent antibodies or was elongated by actin labelled with donor only (F). The average *Δf*_*FRET*_ at the region of 0.3 μm length from the pointed end was estimated from the linear fit.

Furthermore, we examined whether these differences occurred from barbed or pointed ends of the actin filament. To distinguish the barbed end of actin filaments, the ends were capped with gelsolin which was brightly visualised as a large circle by decorating gelsolin-antibody together with a fluorescently labelled secondary antibody (Figure 2E inset). The end without gelsolin signal, presumably the pointed end, was subjected to *Δf*_FRET_ analysis, and all *Δf*_FRET_ from the ten independent experiments belonged to the lower distribution (open triangle in figure 2D inset). To verify, another approach was performed. A donor-labelled actin monomer was added after the filament was formed and decorated with phalloidin. By recognising the elongated end as barbed end (arrow in figure 2F inset), we analysed the *Δf*_FRET_ from the other end, which is likely pointed end. The *Δf*_FRET_ from the ends of all the 10 actin filaments belonged to the distribution with a small peak (black triangle in Figure 2D inset), showing that the small *Δf*_FRET_ is from the pointed end. These results confirmed that the fluctuations in FRET efficiency measured in this study reflect structural polarity along the actin filament.

## 4 Discussion

We used FRET-actin filaments by mixing actin labelled with two different fluorescent dyes. Changes in FRET efficiency showed that actin molecules within the filaments fluctuated over time. In addition, the fluctuation property was different for each end. We first suspected that Brownian motion at the end of the actin filaments might result in random defocusing causing this fluctuation. However, random defocusing should cause the same positional changes at both ends. Our results suggest that *Δf*_*FRET*_ in each end has a different type of fluctuation based on amplitude and decay. The *Δf*_*FRET*_ decreased at a shorter distance from the barbed end to the interior region, compared to the pointed end (Figure 2B). We could not determine how long the *f*_*FRET*_ from both ends affected the interior of the filament because the change in *f*_*FRET*_ from both ends attenuated smoothly as shifted to the interior region of the filaments. Therefore, we analysed the changes in *f*_*FRET*_ in the limited length of 0.2 μm from each end (i.e. ∼70 molecules). This data on *f*_FRET_ difference (Figures 2C and 2D) demonstrated that the intrinsic characteristics must originate from the filament polarity, and the characteristics do not disappear when actin filaments are stabilized with phalloidin.

Our present study showed that the absolute value of FRET efficiency was high in the interior region of actin filaments with lower fluctuations. This high FRET efficiency indicated that the distance change between actin subunits was small. We hypothesized that the small *f*_*FRET*_ can be occurred from following three ways. First, individual actin subunits hardly change their positions. This does not seem probable since our assays were performed in solution and Brownian motion cannot be ignored. The second possible cause could be the random fluctuation of individual subunits and cancelling each other. Randomly changing values of individual molecules were cancelled when they were added (Kozuka et al., 2006). The third possible cause is that fluctuations might cooperatively occur in dozens of actin subunits. We believe that this subunits-cooperativity may also contribute to the formation of a cluster of cofilins extending towards the pointed end, as described in a previous study (Ha et al., 2002; Noguchi et al., 2015). This behaviour suggests that the various states of the actin subunits may be formed at the cluster level, but not with individual subunits and the variation might propagate to the internal regions. When phalloidin binds, cluster-cluster binding is stabilised and fluctuates less, resulting in a smaller *Δf*_*FRET*_.

While this study did not provide any data regarding nucleotide state and structural polymorphism as a possible scenario for the local characteristic change, all-atom simulations presented that actin molecules located near the ends have higher fluctuations than the interior of the actin filaments (Zsolnay et al., 2020). Our data displayed that actin subunits near the terminal ends are constantly fluctuating regardless of the addition of a new actin monomer, suggesting that fluctuations in neighbouring subunits are also sustained. In the simulation by Zsolnay et al., the dihedral angle in the 40th actin molecule from the barbed end was flattened to the same extent as the interior region of the filament, but our *Δf*_*FRET*_ indicated that the effect of the barbed end extended to the inner ∼70th actin. Our data also exhibited that the small *Δf*_FRET_ decreased more gradually toward the interior region at the pointed ends than at the barbed ends. If the adjacent actin subunits fluctuate in the similar orientation, the fluctuation can be coordinated and occurred as the size of clusters. Assuming that the constant number of clusters receives feedback from ends, we think that the cluster size may be larger near the pointed end than the barbed end which may propagate the orientation towards the interior of actin filaments. In this study, we visualised the flexibility of actin filaments using light microscopy. The fluctuation was lower near the end regions than in the interior region. The quantified data showed that the fluctuation was gradually decreased at the pointed end than at the barbed end. Furthermore, the fluctuation decay was more steep at the barbed end than at the pointed end. Once actin filaments are formed, the new orientations, that are governed by the local, spatial and temporal position of actin filaments are generated. Those orientations may take roles to maintain the dynamic change of the actin cytoskeleton.

## 5 Conflict of Interest

The authors declare no conflict of interest.

## 6 Author Contributions

The experimental design was based on HH, IF, TM, and RM. Data correction of the experiments and analyses was performed using TM, MN, and RM. IN and KK signed the program for image analyses. The manuscript was written by TM, HH, and IF. All the authors approved the final version of the manuscript.

## 7 Funding

This study was supported by Grants-in-Aid for Scientific Research C (MEXT KAKENHI) (grant number JP20K06591 to IF), Research Foundation of Opto-Science and Technology to IF and Nagaoka University of Technology Research Grant 2021 for IF.

## 8 Acknowledgements

We thank Ai Takahashi, Kuruto Toda, Hitotaka Ito, and Kazutaka Mori for technical assistance with the sample preparation and microscope setup. We also thank Dr Itsuki Kunita for the image analysis discussion.

## Notes

### Competing Interest Statement

The authors have declared no competing interest.

